# Marine deforestation leads to widespread loss of ecosystem function

**DOI:** 10.1101/852541

**Authors:** Matthew S Edwards, Brenda Konar, Ju-Hyoung Kim, Scott Gabara, Genoa Sullaway, Tristin A McHugh, Michael Spector, Sadie L Small

## Abstract

Trophic interactions can result in changes to the abundance and distribution of habitat-forming species that dramatically reduce ecosystem health and functioning. Nowhere may this be as dramatic as in the coastal zone of the Aleutian Archipelago, where overgrazing by herbivorous sea urchins that began in the 1980s resulted in widespread deforestation of the region’s kelp forests. Here we show that this deforestation resulted in decreased macroalgal and invertebrate abundance and diversity, increased benthic irradiances, and reduced rates of gross primary production and respiration by the ecosystem. These opposing metabolic processes remain in balance, however, which resulted in little-to-no changes to net ecosystem production. These patterns were consistent across nine islands spanning more than 1000 kilometers of the archipelago. In light of the worldwide declines in kelp forests observed in recent decades, our findings suggest that marine deforestation profoundly affects the health of coastal ecosystems and how they function.

**Significance statement:** Widespread marine deforestation results in reduced biodiversity and primary productivity throughout more than 1000 km of the Aleutian Archipelago.

## Introduction

Predators fundamentally affect ecosystems through trophic interactions (*1*). These interactions are especially important if they result in changes to the abundance or distribution of ecosystem engineers, such as forest-forming trees, which can lead to changes in microclimates, biodiversity, primary production, nutrient cycling, and energy flow (*2*). For example, the reintroduction of gray wolves (*Canis lupus*) into Yellowstone National Park, USA in the 1990s resulted in increased predation on elk (*Cervus elaphus*) and subsequently reduced herbivory on canopy-forming trees such as aspens (*Populus tremuloides*), willows (*Salix* spp.), and cottonwoods (*Populus* spp.) (*3*). This ultimately led to changes in the morphology and hydrology of the region’s river systems and its riparian plant communities (*4, 5*). Similarly, large marine algae, such as kelps, can form subtidal forests whose biogenic structures alter hydrodynamic, nutrient and light conditions, modify patterns of biodiversity, enhance primary production and carbon sequestration, and provide food and habitat for numerous other species (*6–9*). Consequently, the loss of these forest-forming kelps and the benthic macroalgae they support can have dramatic impacts to how nearshore ecosystems function, especially if they occur over large geographic areas. Indeed, kelp deforestation has occurred worldwide in recent decades due to a variety of forcing factors (*10, 11*), and the subtidal rocky reefs of the Aleutian Archipelago serve as a model system to investigate the broader impacts of such deforestation. Here, the collapse of sea otter (*Enhydra lutris*) populations led to large increases in their primary prey, herbivorous sea urchins (*Strongylocentrotus polyacanthus*), which subsequently resulted in overgrazing and widespread losses of the region’s kelp forests (*12*). This collapse began in the late 1980s, likely in response to a dietary shift by killer whales toward sea otters, and by 2000 sea otter densities had declined throughout the archipelago to around 5-10% of their estimated equilibrium density (*13*). Currently, most of the kelp forests have either disappeared from the archipelago or are in the process of disappearing, although some small forests remain in their ‘historical state’ at scattered locations on most of the islands (*14, 15*) (Fig. 1). These remnant forests provide an excellent benchmark against which we evaluated the effects of widespread deforestation on two important metrics of ecosystem health and function, namely biodiversity and primary productivity.

**Fig. 1.**
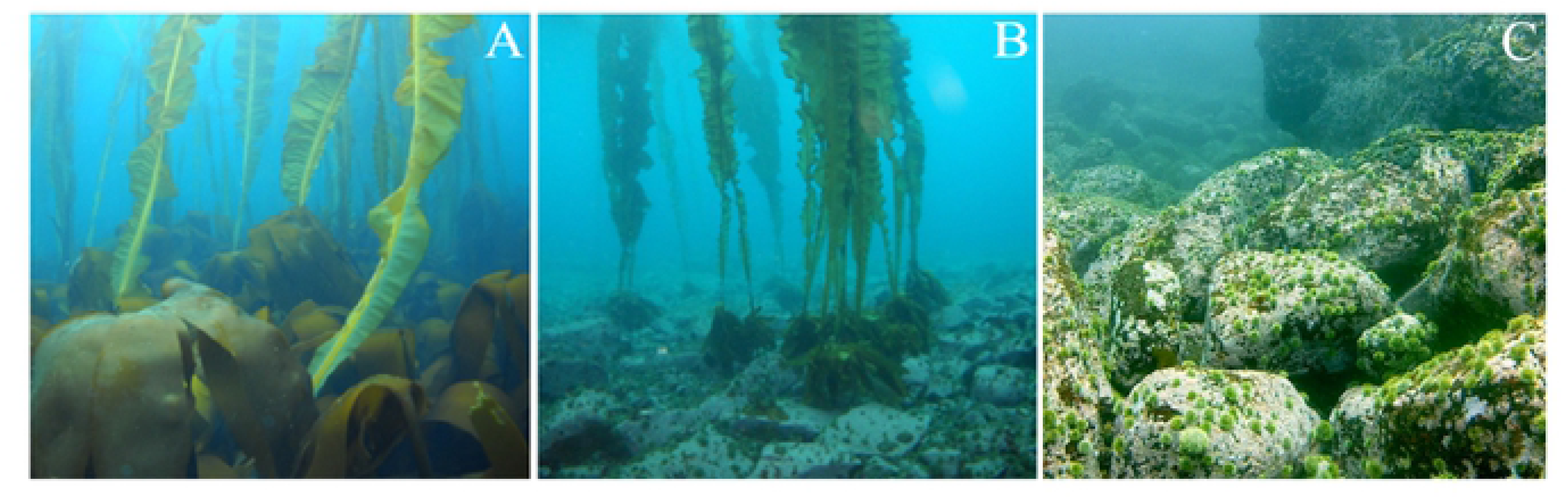
Photographs of each habitat type showing (A) high abundance of benthic macroalgae and canopy-forming kelps in the kelp forests, (B) lack of benthic macroalgae but remaining canopy-forming kelps and high abundances of sea urchins in the transition habitats, and (C) lack of benthic macroalgae and canopy-forming kelps, but high abundances of sea urchins in the urchin barrens.

Characterizing patterns of primary productivity is essential to fully understanding ecosystem health and function (*16, 17*). This includes three basic metrics: gross primary production (*GPP*), which describes all the CO_2_ fixed by the autotrophs during photosynthesis, total ecosystem respiration (*Re*), which describes the release of CO_2_ during the production of energy by autotrophs, heterotrophs, decomposers and microbes, and net ecosystem production (*NEP*), which is the difference between *GPP* and *Re* and describes net changes in the total amount of organic carbon in an ecosystem available for consumption, storage and export to adjacent ecosystems, or nonbiological oxidation to carbon dioxide (*18–21*). In general, ecosystems with high rates of *GPP* also exhibit high rates of *Re*, with the central tendency being that *GPP* and *Re* are in balance (i.e. similar in magnitude) and therefore have median *GPP* / *Re* ratios close to 1.0, and *NEP* values near zero (*21, 22*). Indeed, a review of five decades (1950 to 1990) of studies in aquatic ecosystems demonstrated that these two opposing processes are indeed generally in balance, although unproductive ecosystems tend towards net heterotrophy with *GPP* / *Re* < 1.0 and *NEP* < 0, while productive ecosystems tend towards net autotrophy with *GPP* / *Re* > 1.0 and *NEP* > 0 (*19–22*). Further, the amount of *Re* associated with any given *GPP* in shallow coastal ecosystems tends to be greater when the complete benthic communities are considered (*22*). This may be especially true if microbial metabolism, which is an important component of *Re*, is large compared to *GPP* (*19–21*). This is important for coastal kelp forests, which generally have higher microbial diversity relative to the adjacent ocean waters (*23–25*). Consequently, loss of these forests may lead to complex patterns of *GPP, Re* and *NEP* within coastal ecosystems. On one hand, reductions in primary producer biomass should result in lowered *GPP* and thus reduced *NEP*. Alternately, deforestation may result in reduced biodiversity and lowered abundances of macroalgae, invertebrates, fishes and microbes, which may lead to reduced *Re* and enhanced *NEP*. At the same time, loss of the habitat-forming kelps also results in increased benthic irradiances (*17*) and thus potentially to increased compensatory production by any remaining fleshy macroalgae, encrusting coralline algae, and microalgae (*26–28*), which can result in enhanced *NEP*. Thus, understanding the balance between *GPP* and *Re* in addition to *NEP* can be instrumental in discerning the broader impacts of deforestation on ecosystem health and productivity. This may be especially relevant for the Aleutian Archipelago where widespread kelp deforestation has resulted in significant reductions in fishes, invertebrates and fleshy macroalgae, increases in encrusting coralline algae (*12, 29*), and elevated benthic irradiances (*14*).

## Results

We studied patterns of benthic macroalgal and invertebrate abundance and diversity, and rates of *NEP, GPP* and *Re* within remnant kelp forests, urchin barrens, and habitats that were in transition to becoming urchin barrens (Fig. 1) at nine islands spanning more than 1000 kilometers of the Aleutian Archipelago (Fig. 2, Table 1). These kelp forest and urchin barrens occur as alternate stable states of one another, often with sharply delineated boundaries between them (*15*). Our results show that the benthic communities within the remnant kelp forests have more than a 10-fold greater biomass of fleshy macroalgae than those in the urchin barrens (Permutation post hoc: p = 0.004), while the urchin barrens have a nearly 3-fold greater biomass of urchins than the kelp forests (Fisher’s LSD: p < 0.001, Fig. 3, Tables 2 & 3). The kelp forests also had greater diversity of macroalgae and encrusting invertebrates than either the urchin barrens or transition habitats (*30*), primarily due to the presence of large kelps within the kelp forests and abundant sea urchins within the barren grounds (Fig. 4). The transition habitats were similar to (i.e. did not differ from) the urchin barrens with high abundances of urchins (p = 0.096) and low abundance of fleshy macroalgae (p = 0.120) on the benthos (Fig. 3), and are similar to the kelp forests in the mid-water and at the surface with many canopy-forming kelp (*Eualaria fistulosa*) still remaining (Fig. 1). All three habitats have high bottom covers of encrusting coralline algae that lie below the fleshy macroalgae and become exposed following deforestation (Fig. 1). Benthic irradiances (*PAR – photosynthetically active radiation*) vary among the three habitat types (ANOVA: F_2,16_ = 7.697, p = 0.004) and are greatest in the urchin barrens, lowest in the kelp forests, and intermediate in the transition habitats (Fig. 3, Tables 4 & 5).

**Table 1.** List of the nine islands in the Aleutian Archipelago where we deployed cBITs to measure *NEP, GPP* and *Re,* and the six islands where we collected all macroalgae and invertebrates from within the cBITs to estimate their biomass. The number of cBITs deployed, and whether macroalgae and invertebrates were collected from within them are noted.

| <u>Island</u> | <u>No. cBITs deployed</u> |  |  | <u>Collections made?</u> |
| --- | --- | --- | --- | --- |
|  | Kelp<br>forests | Transition<br>habitats | Urchin<br>barrens |  |
| Adak | 3 | 2 | 3 | No |
| Amchitka | 3 | 2 | 3 | Yes |
| Atka | 3 | 3 | 2 | Yes |
| Attu | 3 | 3 | 1 | Yes |
| Chuginadak | 2 | 2 | 3 | Yes |
| Kiska | 3 | 3 | 2 | Yes |
| Nizki | 3 | 3 | 2 | Yes |
| Tanaga | 2 | 1 | 2 | No |
| Yunaska | 3 | 3 | 3 | No |

**Table 2.** Results of A) a two-way Model III Permutation Analysis of Variance testing differences in Algae biomass, and B) a two-way Model III Analyses of Variance testing differences in Urchin biomass, among the nine islands and and three habitat types (kelp forests, transition habitats, and urchin barrens). For each analysis, island was considered a random factor and habitat was considered a fixed factor.

**A) Algae biomass**
| Source | Type III SS | df | MS | Pseudo-F | p-value |
| --- | --- | --- | --- | --- | --- |
| Island | 5 | 4.533 | 0.907 | 6.146 | 0.001 |
| Habitat | 2 | 32.126 | 16.063 | 24.525 | 0.002 |
| Hab*Isl | 10 | 6.550 | 0.655 | 4.440 | 0.002 |
| Error | 36 | 5.310 | 0.148 |  |  |

**B) Urchin biomass**
| Source | Type III SS | df | MS | F-ratio | p-value |
| --- | --- | --- | --- | --- | --- |
| Island | 0.837 | 5 | 0.167 | 3.523 | 0.011 |
| Habitat | 4.185 | 2 | 2.092 | 10.676 | 0.003 |
| Hab*Isl | 1.962 | 10 | 0.196 | 4.131 | 0.001 |
| Error | 1.71 | 36 | 0.047 |  |  |

**Table 3.** Results of A) permutation post hoc comparisons testing for differences in macroalgal biomass, and B) Fisher’s LSD pairwise comparisons testing for differences in urchin biomass, among habitat type pairs. These tests were done as *a priori* hypotheses and thus done regardless of PERMANOVA or ANOVA outcomes (see Table 2).

**A) Macroalgae**
| Habitat 1 | Habitat 2 | t | P(perm) | perms |
| --- | --- | --- | --- | --- |
| Barren | Kelp | 12.266 | 0.004 | 960 |
| Barren | Transition | 1.811 | 0.120 | 974 |
| Kelp | Transition | 3.991 | 0.020 | 974 |

**B) Urchins**
| Habitat 1 | Habitat 2 | Difference | p-value | 95.0% Confidence Intervals |  |
| --- | --- | --- | --- | --- | --- |
|  |  |  |  | Lower | Upper |
| Barren | Kelp | 0.643 | <0.001 | 0.495 | 0.79 |
| Barren | Transition | 0.124 | 0.096 | -0.023 | 0.272 |
| Kelp | Transition | -0.518 | <0.001 | -0.666 | -0.371 |

**Table 4.** Results of separate two-way Model III Analyses of Variance testing for differences in A) net ecosystem production (*NEP*), B) gross primary production (*GPP*), C) respiration (*Re*), D) the range (difference) between *GPP* and *Re*, and E) *PAR* among the nine islands and three habitats (kelp forests, transition habitats, and urchin barrens). For each analysis, island was considered a random factor and habitat was considered a fixed factor.

| **A) *NPP*** |  |  |  |  |  |
| --- | --- | --- | --- | --- | --- |
| Source | Type III<br>SS | df | MS | F-ratio | p-value |
| Island | 7.98E+02 | 8 | 99.783 | 4.623 | 0.001 |
| Habitat | 93.089648 | 2 | 46.545 | 0.502 | 0.314 |
| Hab*Isl | 1.48E+03 | 16 | 92.561 | 4.289 | 0.001 |
| Error | 8.85E+02 | 41 | 21.583 |  |  |
| **B) *GPP*** |  |  |  |  |  |
| Source | Type III<br>SS | df | MS | F-ratio | p-value |
| Island | 4.57E+03 | 8 | 5.71E+02 | 8.077 | 0.001 |
| Habitat | 2.65E+02 | 2 | 1.33E+02 | 0.788 | 0.471 |
| Hab*Isl | 2.69E+03 | 16 | 1.68E+02 | 2.378 | 0.013 |
| Error | 2.90E+03 | 41 | 7.07E+01 |  |  |
| **C) *Re*** |  |  |  |  |  |
| Source | Type III<br>SS | df | MS | F-ratio | p-value |
| Island | 4.58E+03 | 8 | 5.73E+02 | 9.766 | 0.001 |
| Habitat | 4.94E+02 | 2 | 2.47E+02 | 1.246 | 0.314 |
| Hab*Isl | 3.17E+03 | 16 | 1.98E+02 | 3.375 | 0.001 |
| Error | 2.41E+03 | 41 | 58.684 |  |  |
| **D) Range** |  |  |  |  |  |
| Source | Type III<br>SS | df | MS | F-ratio | p-value |
| Island | 8.77E+03 | 8 | 1.10E+03 | 8.857 | 0.001 |
| Habitat | 7.36E+02 | 2 | 3.68E+02 | 1.077 | 0.363 |
| Hab*Isl | 5.46E+03 | 16 | 3.41E+02 | 2.758 | 0.005 |
| Error | 5.07E+03 | 41 | 1.24E+02 |  |  |
| **E) *PAR*** |  |  |  |  |  |
| Source | Type III<br>SS | df | MS | Pseudo-F | p-value |
| Island | 11.856 | 2 | 5.928 | 7.964 | 0.004 |
| Habitat | 6.074 | 8 | 0.759 | 1.020 | <0.001 |
| Hab*Isl | 11.909 | 16 | 0.744 | 6.554 | <0.001 |
| Error | 3.748 | 33 | 0.114 |  |  |

**Table 5.** Results of Fisher’s LSD pairwise comparisons testing for differences in A) *NEP,* B) *GPP,* C) *Re,* D) the range (difference) between *GPP* and *Re,* and E) *PAR* among habitat type pairs. These tests were carried out as *a priori* hypotheses, and thus done regardless of ANOVA outcomes (see Table 4).

| **A) *NEP*** |  |  |  |  |  |
| --- | --- | --- | --- | --- | --- |
| Habitat 1 | Habitat 2 | Difference | p-value | 95.0% Confidence Intervals |  |
|  |  |  |  | Lower | Upper |
| Barren | Kelp | 1.289 | 0.642 | -2.055 | 4.633 |
| Barren | Transition | -1.647 | 0.523 | -5.093 | 1.800 |
| Kelp | Transition | -2.936 | 0.107 | -6.238 | 0.367 |

| **B) *GPP*** |  |  |  |  |  |
| --- | --- | --- | --- | --- | --- |
| Habitat 1 | Habitat 2 | Difference | p-value | 95.0% Confidence Intervals |  |
|  |  |  |  | Lower | Upper |
| Barren | Kelp | -4.871 | 0.067 | -9.898 | 0.155 |
| Barren | Transition | -1.719 | 0.532 | -6.899 | 3.462 |
| Kelp | Transition | 3.153 | 0.225 | -1.811 | 8.117 |

| **C) *Re*** |  |  |  |  |  |
| --- | --- | --- | --- | --- | --- |
| Habitat 1 | Habitat 2 | Difference | p-value | 95.0% Confidence Intervals |  |
|  |  |  |  | Lower | Upper |
| Barren | Kelp | -6.311 | 0.011 | -10.890 | -1.731 |
| Barren | Transition | -1.235 | 0.621 | -5.955 | 3.485 |
| Kelp | Transition | 5.076 | 0.035 | 0.553 | 9.598 |

| **D) RANGE** |  |  |  |  |  |
| --- | --- | --- | --- | --- | --- |
| Habitat 1 | Habitat 2 | Difference | p-value | 95.0% Confidence Intervals |  |
|  |  |  |  | Lower | Upper |
| Barren | Kelp | -7.838 | 0.027 | -14.487 | -1.188 |
| Barren | Transition | -1.887 | 0.603 | -8.740 | 4.966 |
| Kelp | Transition | 5.951 | 0.086 | -0.616 | 12.517 |

| **E) *PAR*** |  |  |  |  |  |
| --- | --- | --- | --- | --- | --- |
| Habitat 1 | Habitat 2 | Difference | p-value | 95.0% Confidence Intervals |  |
|  |  |  |  | Lower | Upper |
| Barren | Kelp | 1.090 | <0.001 | 0.876 | 1.304 |
| Barren | Transition | 0.456 | <0.001 | 0.239 | 0.673 |
| Kelp | Transition | -0.634 | <0.001 | -0.854 | -0.415 |

**Fig. 2.**
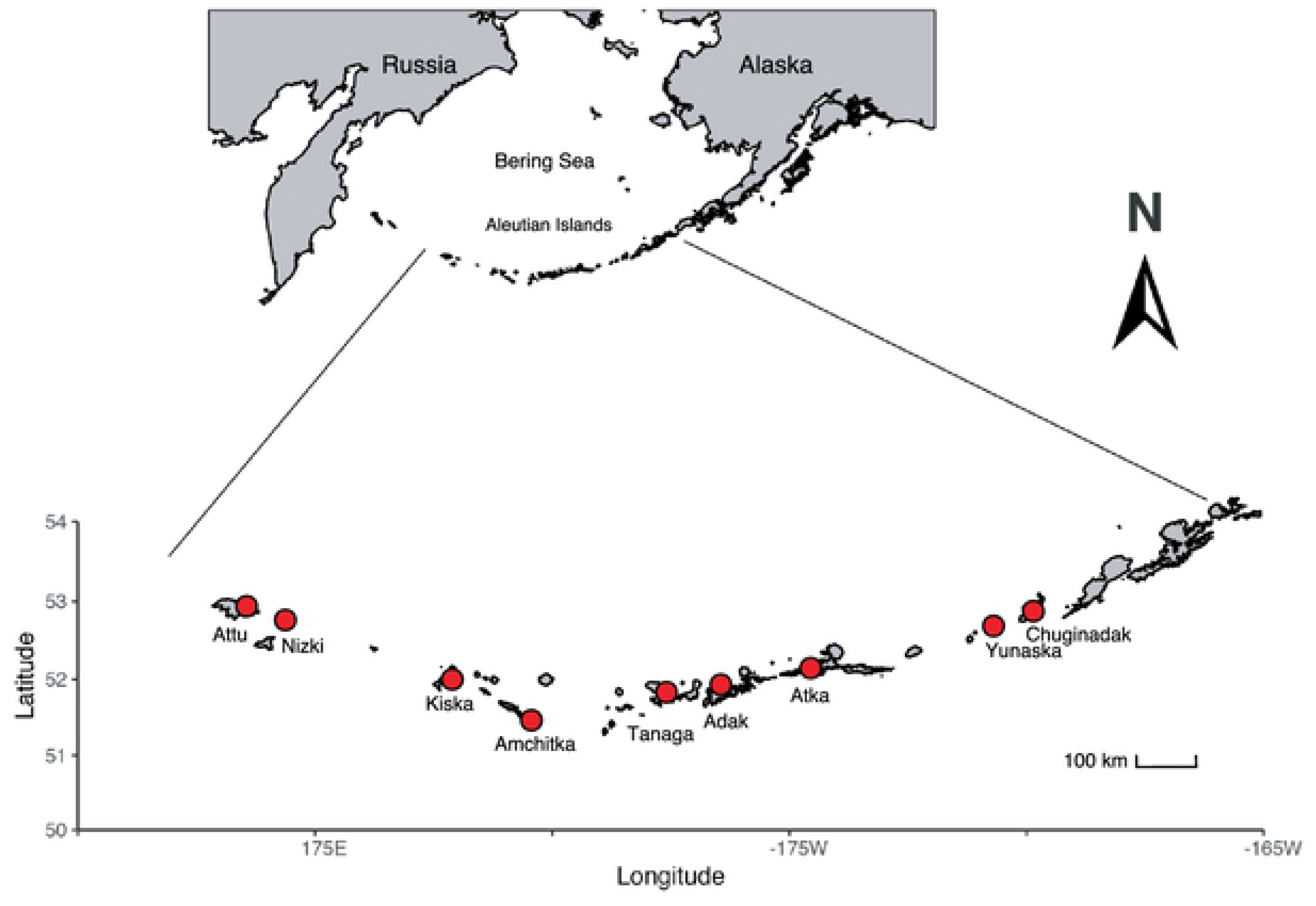
Map of the Aleutian Archipelago showing locations of the nine islands (denoted by red circles) where ecosystem productivity (*NEP*, *GPP* and *Re*) was measured in the cBITs.

**Fig. 3.**
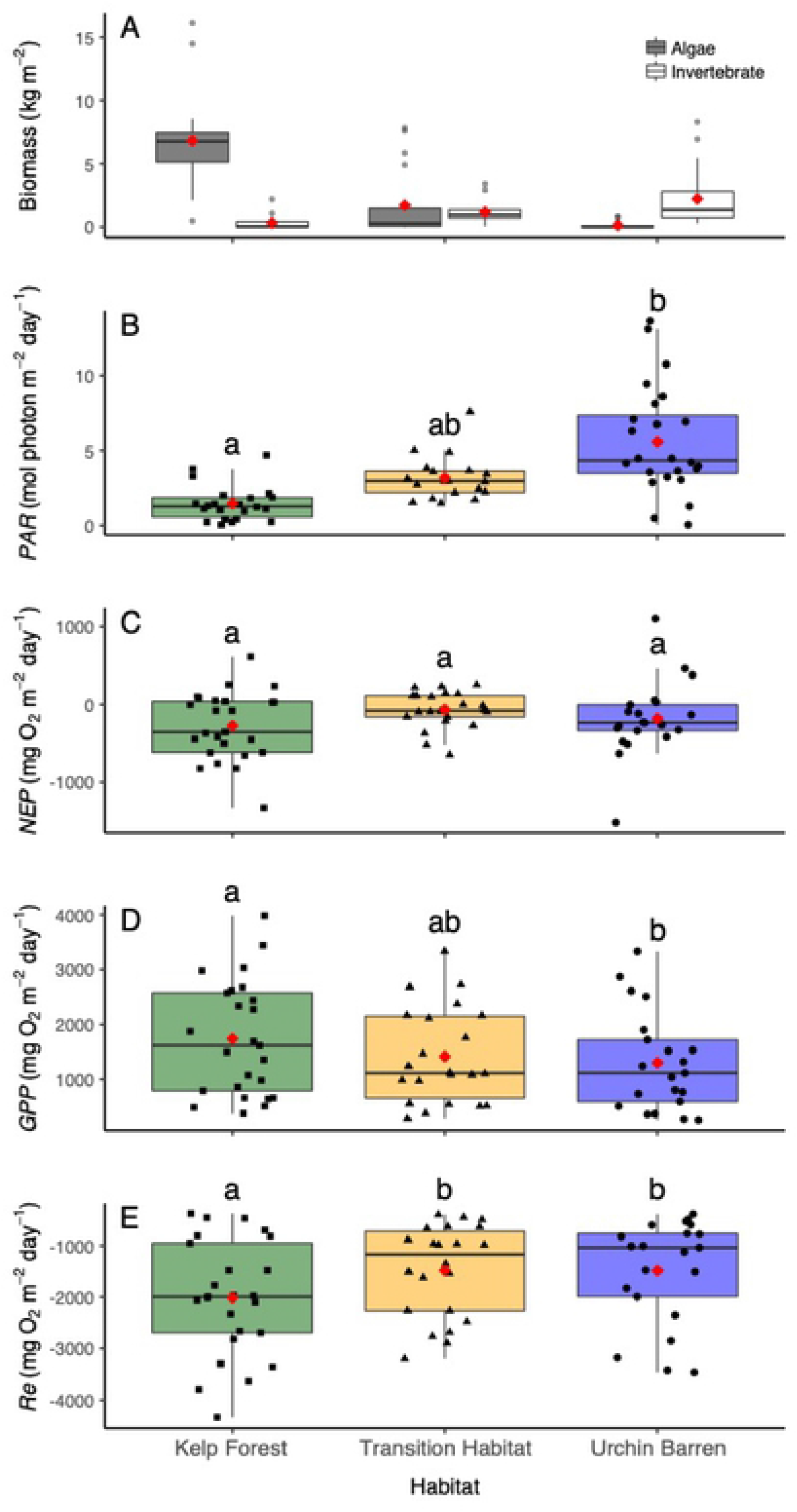
Box plots showing (A) Macroalgae (gray bars) and invertebrate (white bars) biomass, (B) Irradiance (*PAR*), (C) Net Ecosystem Production (*NEP*), (D) Gross Primary Production (*GPP*), and (E) Ecosystem Respiration (*Re*), as measured in the cBITs deployed within each habitat type (kelp forests, transition habitats, and urchin barrens). Macroalgae and invertebrate diversity and biomass were measured at six islands, and *PAR*, *GPP*, *Re*, and *NEP* were measured at nine islands (Fig 2, Table 1). Red diamonds represent mean values, and horizontal lines represent median values. Boxes within each graph that do not share letters represent significant differences between habitat pairs.

**Fig. 4.**
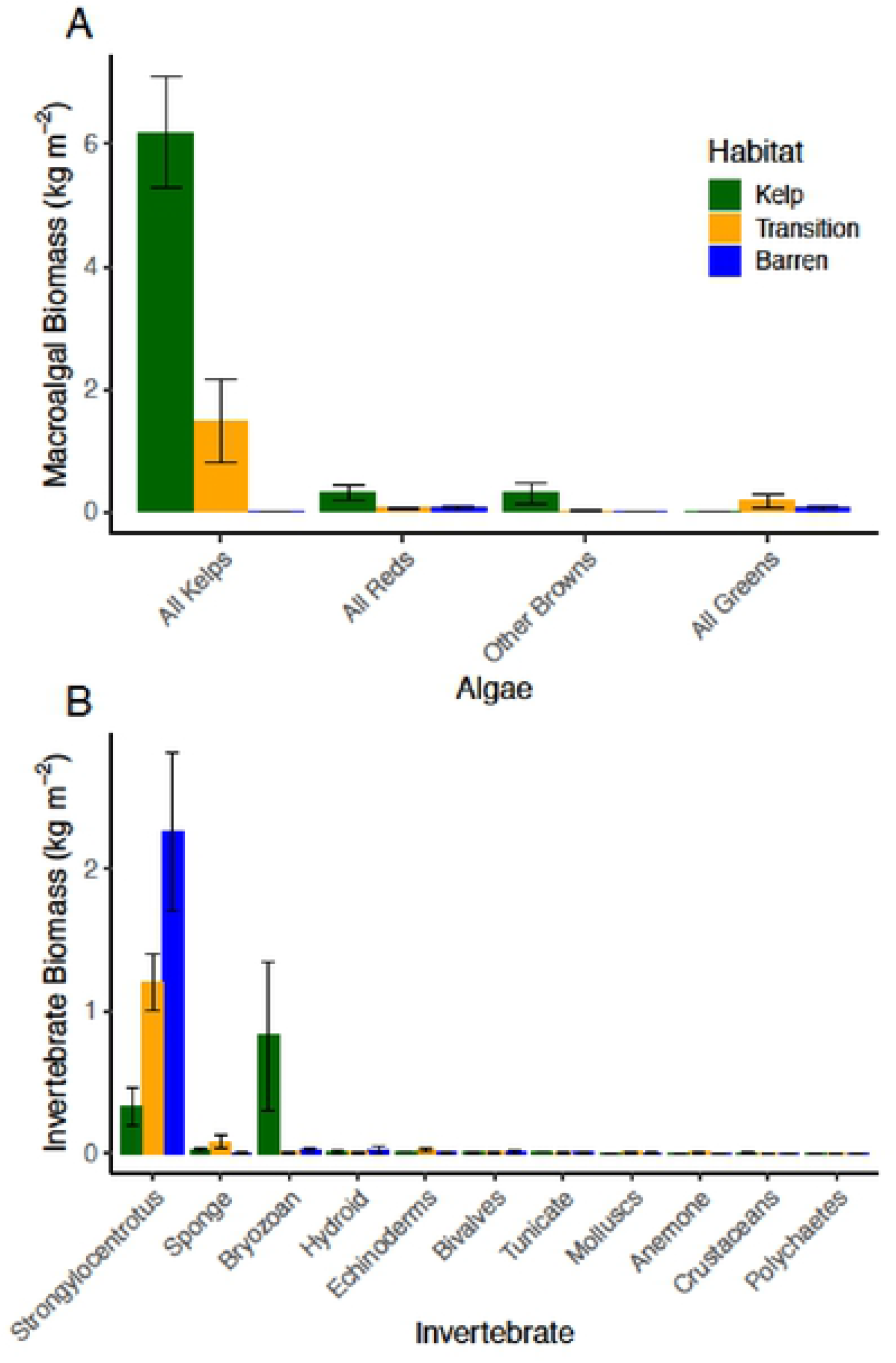
Mean biomass (± SE) of (A) all kelps, and red, brown and green macroalgae, and (B) the most abundant taxonomic groups of invertebrates collected within the cBITs within each habitat type at six of the islands where the cBITs were deployed (Table 1).

We examined how the differences in benthic communities and *PAR* influenced *NEP*, *GPP, Re* and the balance between *GPP* and *Re* by measuring changes in seawater oxygen concentrations within replicate (n = 3) chambers (collapsible benthic incubation tents; hereafter cBITs) that were placed on the benthos over representative assemblages within each habitat type at each island. We predicted that *NEP* at the benthos would be reduced in the urchin barrens due to the loss of photosynthetic macroalgae. Instead, we found that *NEP* does not differ between any of the habitat types, nor does it differ from zero (i.e., *GPP* = *Re*) in any of the habitat types (Figs. 3 & 5; Tables 4 & 5). Benthic *GPP* in contrast, was 33% higher in the kelp forests than in the urchin barrens (Fisher’s LSD: p = 0.067), and 23% higher in the kelp forests than in the transition habitats (p = 0.225), but it differs by only 7% between the transition habitats and urchin barrens (p = 0.532) (Fig. 3, Table 2 & 3). This is presumably due to the higher abundance of benthic fleshy macroalgae in the kelp forests, but similar abundances of fleshy macroalgae in the urchin barrens and transition habitats (Fig. 4). Similarly, benthic *Re* is 35% higher in the kelp forests than it is in both the urchin barrens (Fisher’s LSD: p = 0.011) and the transition habitats (p = 0.035), but it differs by less than 1% between the transition habitats and the urchin barrens (p = 0.621) (Fig. 3, Table 4 & 5). This is presumably due to the higher biomass of fleshy macroalgae and invertebrates, lower irradiances, and greater diversity of kelp-associated microbes (*23–25*) in the kelp forests, while the urchin barrens and transition habitats have similarly high abundances of urchins and low biomasses of macroalgae. Lastly, the difference (i.e. range) between *GPP* and *Re*, which we believe to be a better measure of ecosystem function than *NEP*, is 34% greater in the kelp forests than in the urchin barrens (Fisher’s LSD: p = 0.027), and 29% greater in the kelp forests than in the transition habitats (p = 0.086), but this range varies by less than 4% between the transition habitats and the urchin barrens (p = 0.603) (Fig. 3, Table 4 & 5). Thus, while we expected *NEP* to scale positively with autotroph biomass by habitat, we found no differences in benthic *NEP* among the three habitat types. Instead, we found that kelp forests have the highest *GPP* and *Re,* and that the urchin barrens and the transition habitats do not differ with respect to these metrics. PAR did vary significantly among the three habitat types and was greater in the urchin barren grounds than in the kelp forests or the transition habitats (Fisher’s LSD: p < 0.001) (Tables 4 & 5). This indicted deforestation resulted in widespread losses to primary production and respiration by the ecosystem, and increases in benthic irradiances.

**Fig 5.**
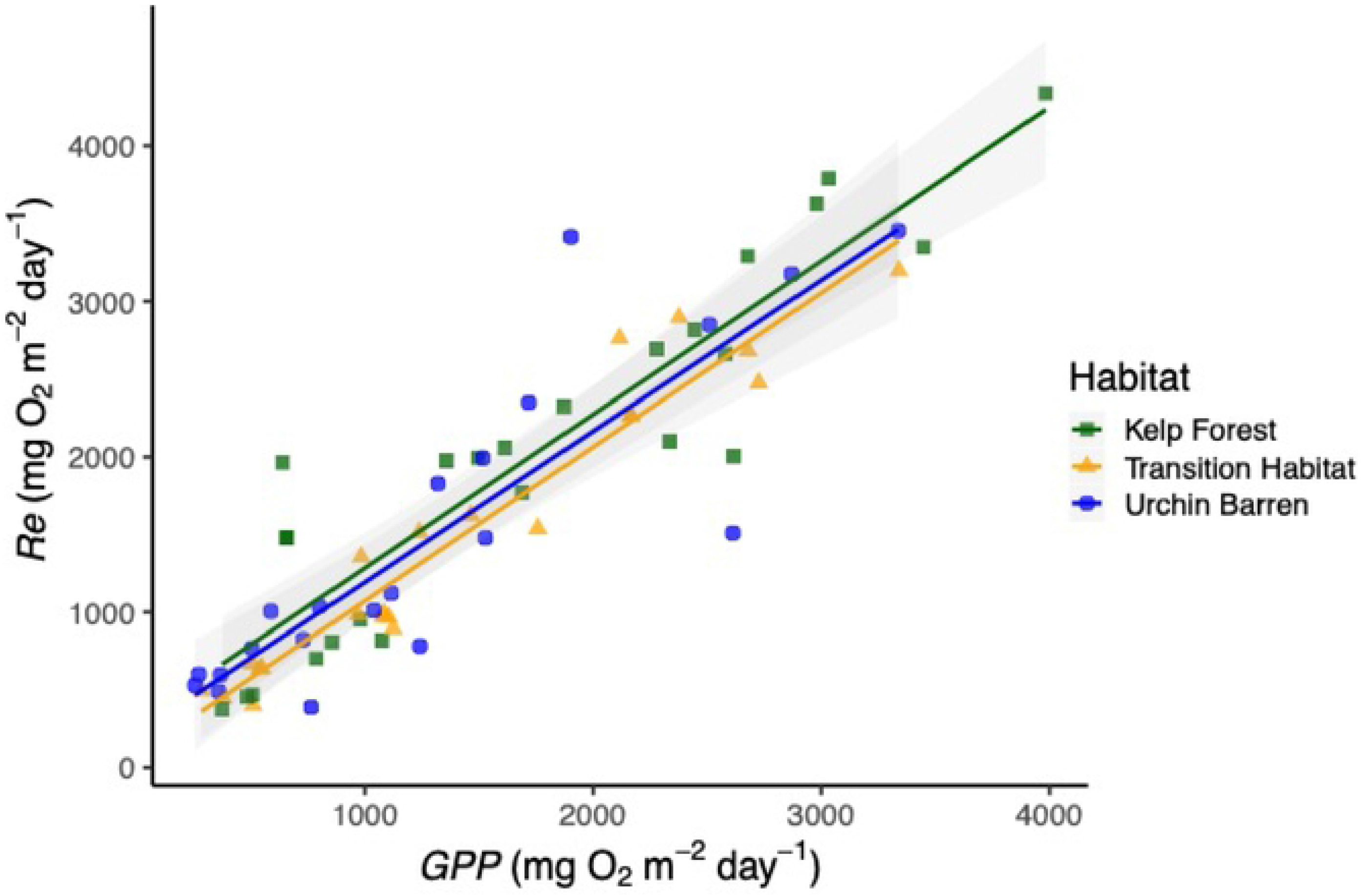
Relationship between gross primary production (*GPP*) and ecosystem respiration (*Re*) for each habitat type across all nine islands where cBITs were deployed (Table S1). Each point represents measurements from a single cBIT. Gray shading denoted 95% confidence intervals.

Our study is in agreement with previous studies in aquatic ecosystems that have shown *GPP* and *Re* to generally be in balance and thus exhibit *GPP* / *Re* ratios near 1.0, and *NEP* values near zero (*21, 22*). Indeed, when we examine the relationships between *GPP* and *Re* in each of the cBITs in each habitat type separately, *GPP* and *Re* are consistently similar in magnitude, with no differences in *GPP / Re* ratios among habitat types (ANCOVA: F_2,62_ = 0.16, p = 0.852) (Fig. 5, Table 6). Further, the distribution of these ratios is symmetrical around 1.0 in each habitat (Fig. 6). Interestingly, the highest individual values of *NEP* were not observed in the kelp forests but rather in the urchin barrens, which we believe was due to higher irradiances in the urchin barrens than the other two habitats (Fig. 3) combined with compensatory production by the encrusting coralline algae and benthic diatoms (*28*). However, those few observations aside, it is clear that all three benthic habitats remain in balance following deforestation, with *GPP* ≈ *Re, GPP / Re* ratios ≈ 1, and median *NEP* values ≈ 0. Thus, although *NEP* may help differentiate between productive and unproductive ecosystems (*22*), it poorly describes changes in primary productivity following widespread kelp deforestation in the Aleutian Archipelago. Instead, it is clear that deforestation results in significant changes to the region’s benthic communities, and these led to declines in both *GPP* and *Re,* which better reflect a reduction in ecosystem functioning (*16, 17*). Further, it appears that even partial deforestation, where the benthic macroalgae and invertebrates have been lost but the canopy-forming kelps remain, results in a decrease in *GPP* and *Re* at the benthos that is similar to trends found in urchin barrens.

**Fig 6.**
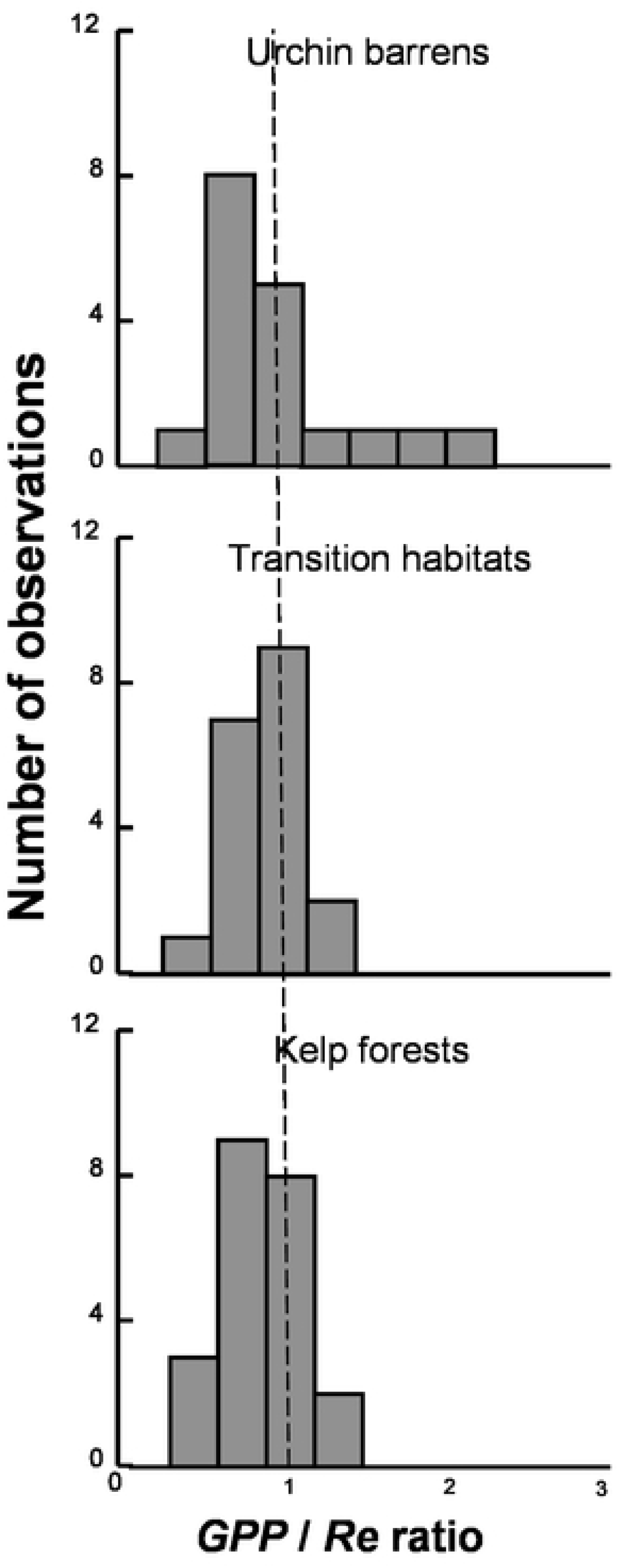
Frequency distribution of *GPP* / *Re* ratios within each habitat type across all nine islands where cBITs were deployed (Table S1). Each data point represents measurements from a single cBIT. Note the urchin barrens have the highest ratios observed, and the kelp forests have the largest number of low values. The vertical dashed line represents the 1:1 ratio.

**Table 6.**
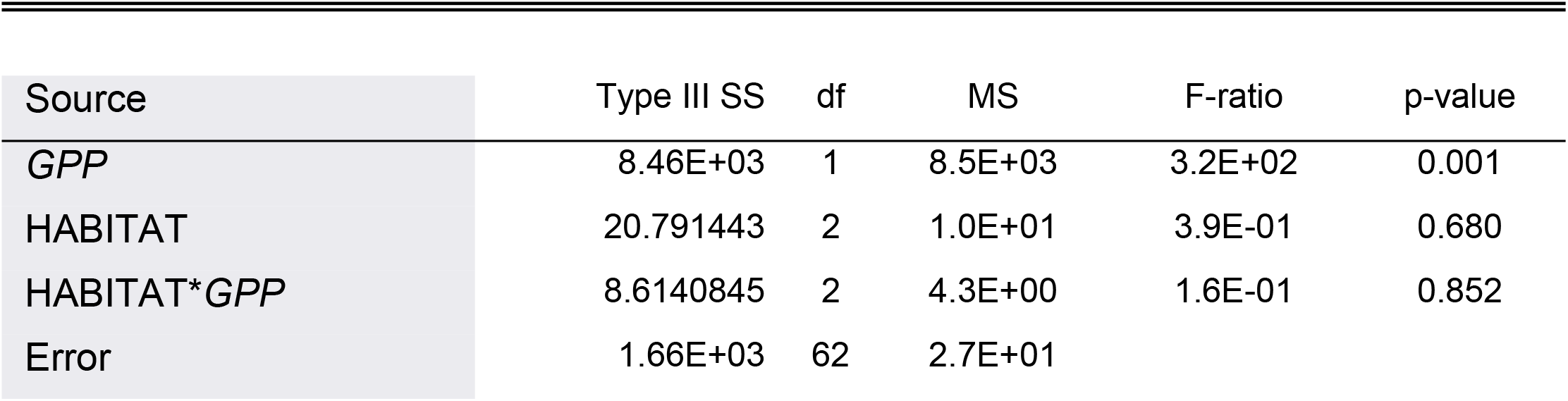
Analysis of covariance testing the effect of *GPP* and habitat on *Re.* Note the non-significant Habitat*\*GPP* interaction hat shows no differences in the slopes (i.e. relationships) between *GPP* and *Re* among the three habitat types. See Fig 5 for graphical representation.

## Discussion

Trophic interactions can lead to changes to the distribution and abundance of habitat-forming species, which can have profound impacts on ecosystem health and function (*2, 31*). Deforestation, in particular, can result in changes to biodiversity and energy flow (*2*), altered regional and global climates (*32*), and even lead to species extinctions (*33*). Coastal kelps are an excellent example of such ecosystem engineers in nearshore habitats that have suffered large-scale deforestation over the past few decades due to both biological and physical stressors (*10, 11*). Consequently, our study is relevant to other areas of the world where kelp forests have exhibited local to broad scale declines, such as the northwest coast of the United States (*34*), Nova Scotia (*35*), western Europe (*36*), southwestern Japan (*37*), the east coast of South Korea (*38*), and along the southern coast of Australia (*39*). Indeed, recent estimates suggest that global declines in kelp abundances may be as high as 2% per year (*11*), which can negatively impact numerous other species that depend on them for food and habitat. Certainly, the kelp forests of the Aleutian Archipelago are in critical condition in the face of widespread overgrazing by urchins, and this has had profound effects on the region’s benthic communities and on patterns of gross primary production and ecosystem respiration. Whether these forests will recover and return to prior ecosystem functioning regarding these metrics is unknown, but observations of kelp forests from other areas of the world suggest it is possible. For example, *Laminaria longicruris* forests recovered from overgrazing following localized disease outbreaks that decimated sea urchin populations in Nova Scotia (*40*), while *L. hyperborea* forests recovered in mid-Norway due to low sea urchin recruitment (*41*). *Ecklonia maxima* expanded its range eastward in South Africa, coincident with cooling of the local ocean waters (*42*). Likewise, *Macrocystis pyrifera* recovered along a ∼100 km stretch of the Pacific coast of Baja California, Mexico following nearly two decades of absence after the strong 1997-98 El Niño Southern Oscillation (*43*). Recovery of the *Eualaria fistulosa* forests throughout the Aleutian Archipelago, however, would likely require widespread mortality in the urchin populations, which today seems unlikely. Until then, benthic biodiversity, *GPP* and *Re* will likely remain lower in areas of kelp forest loss because the high abundance of urchins limits regrowth of macroalgae and maintains the urchin barrens (*15*). Thus, we present a benchmark against which we can evaluate this recovery, and understand the effects of further deforestation in this ecosystem.

Although we have learned much about the effects of the otter-urchin-kelp trophic cascade in the Aleutian Archipelago, this study offers new insights into the consequences of such widespread deforestation on the region’s benthic primary productivity. Certainly, benthic *GPP, Re* and the difference between them are all greatest in the kelp forests where macroalgae, fish, invertebrate, and presumably microbial, communities are all most abundant. Deforestation then resulted in reductions in these metrics, identifying loss of ecosystem health and function regarding biodiversity, macroalgal abundances, and primary productivity. In contrast, benthic biodiversity, macroalgal abundances, *GPP*, *Re* and the difference between them are all similar in the urchin barrens and transition habitats, suggesting that the transition habitats have already suffered reduced ecosystem function following losses of their benthic communities. This, of course, reflects productivity at the benthos and not in the mid-water or at the surface where the canopy-forming *Eualaria fistulosa* remains in the transition habitats. It is likely that these canopy-forming macroalgae would increase *GPP* and perhaps result in positive values of *NEP* in the mid-water and at the surface in both the kelp forests and transition habitats. However, at the benthos, *GPP* and *Re* remain in balance following deforestation, leading to similar, near-zero *NEP* in all three habitats. We believe this reflects balance between the autotrophic and heterotrophic components of the ecosystem. Specifically, the macroalgae exhibit positive *GPP* as they photosynthesize, grow and increase in abundance, but this results in a concomitant increase in heterotrophic metabolism, which increases *Re*. In the face of deforestation, both *GPP* and *Re* decrease, resulting in little to no changes in *NEP*. Thus, we propose that *GPP* and *Re* are better measures of changes to primary productivity than *NEP.* Combining these with estimates of macroalgal and invertebrate diversity and abundance revealed that the Aleutian Archipelago suffered substantial losses to ecosystem function following widespread deforestation.

## Materials and Methods

While many past experiments examining primary production by autotrophic communities have relied on laboratory experiments that do not incorporate natural fluctuations in abiotic conditions, recent studies have identified techniques that measure primary production *in situ*, thereby increasing the ecological realism of their experiments (*44–46*). For example, *in situ* chamber designs have been developed for estimating primary production by individual species (*45, 46*) and whole benthic communities (*27, 46*). In general, estimates of net ecosystem production (*NEP*) on the benthos can be made by measuring changes in dissolved oxygen within benthic chambers that are placed *in situ* over of macroalgae and invertebrate communities. In this study, we deployed collapsible benthic isolation tents (cBITs) modelled after those described by Haas et al. (*47*) and Calhoun et al. (*48*) that directly measured *in situ* benthic oxygen production and allowed us to estimate gross primary production (*GPP*), ecosystem respiration (*Re*) and net ecosystem production *NEP* by the benthic communities (*27, 28, 45*). By linking temporal changes in oxygen concentrations within the cBITs to incident irradiance conditions and organism abundances, we can relate variation in *GPP*, *Re,* and *NEP* to primary producer and invertebrate biomass (*27, 49*). Further, because our cBITs encompassed whole benthic communities, species interactions (e.g., shading), and invertebrate and microbial respiration were incorporated into production measurements. These interactions are often not captured in laboratory experiments but are pertinent to understanding *GPP*, *Re,* and *NEP* (*50*).

### Experimental Design

Our cBITs were made from 0.106 cm polycarbonate plastic triangle sheets glued to fiberglass-reinforced vinyl panels (Fig. 7). The frames were reinforced using stainless steel tubes with stainless steel cable to facilitate handling and to ensure they held their pyramidal shape with an internal volume of 192 L and a basal area of (0.64 m^2^). The cBITs each had 26” skirts around the perimeter, upon which chain was laid to hold them to the benthos and prevent water exchange with the surrounding environment. The polycarbonate walls were thin and flexible to allow hydrodynamic energy transfer into the cBITs, thereby reducing boundary layer formation around the macroalgal thalli. We verified this energy transfer using dissolving plaster blocks placed within cBITs and by using video analysis of internal seaweed movements. Sensor arrays that included a Photosynthetic Active Radiation (*PAR*) sensor (Odyssey Dataflow Systems Ltd), and a Dissolved Oxygen (DO mg/L) and Temperature (°C) sensor (MiniDOT Logger, PME) were placed at the center of each cBIT (Fig 8).

**Fig. 7.**
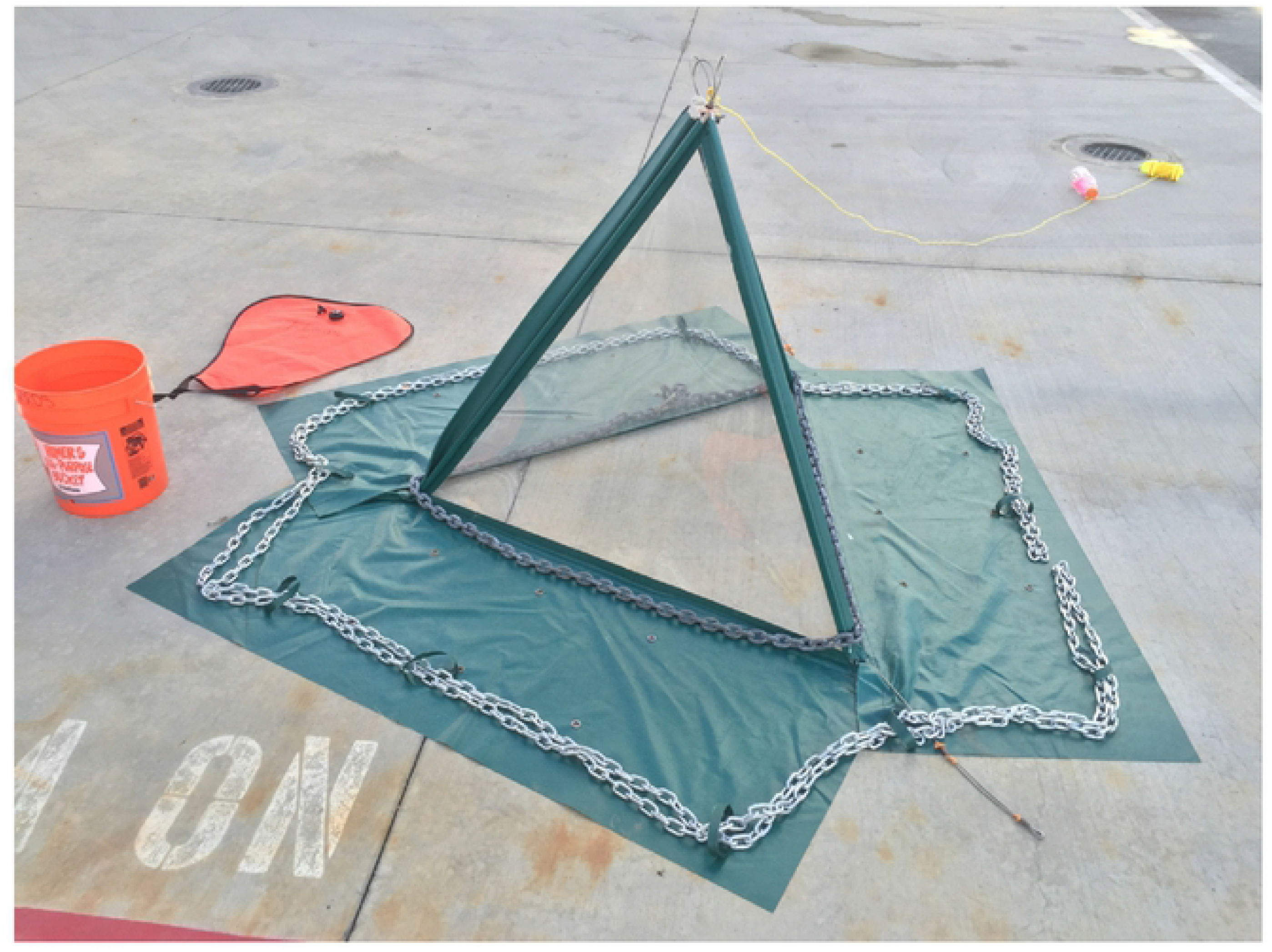
Photograph of cBIT before deployment showing 26” skirt around perimeter, flexible polycarbonate walls, steel framing, anchor chain used to hold skirt and cBIT to the benthos.

**Fig 8.**
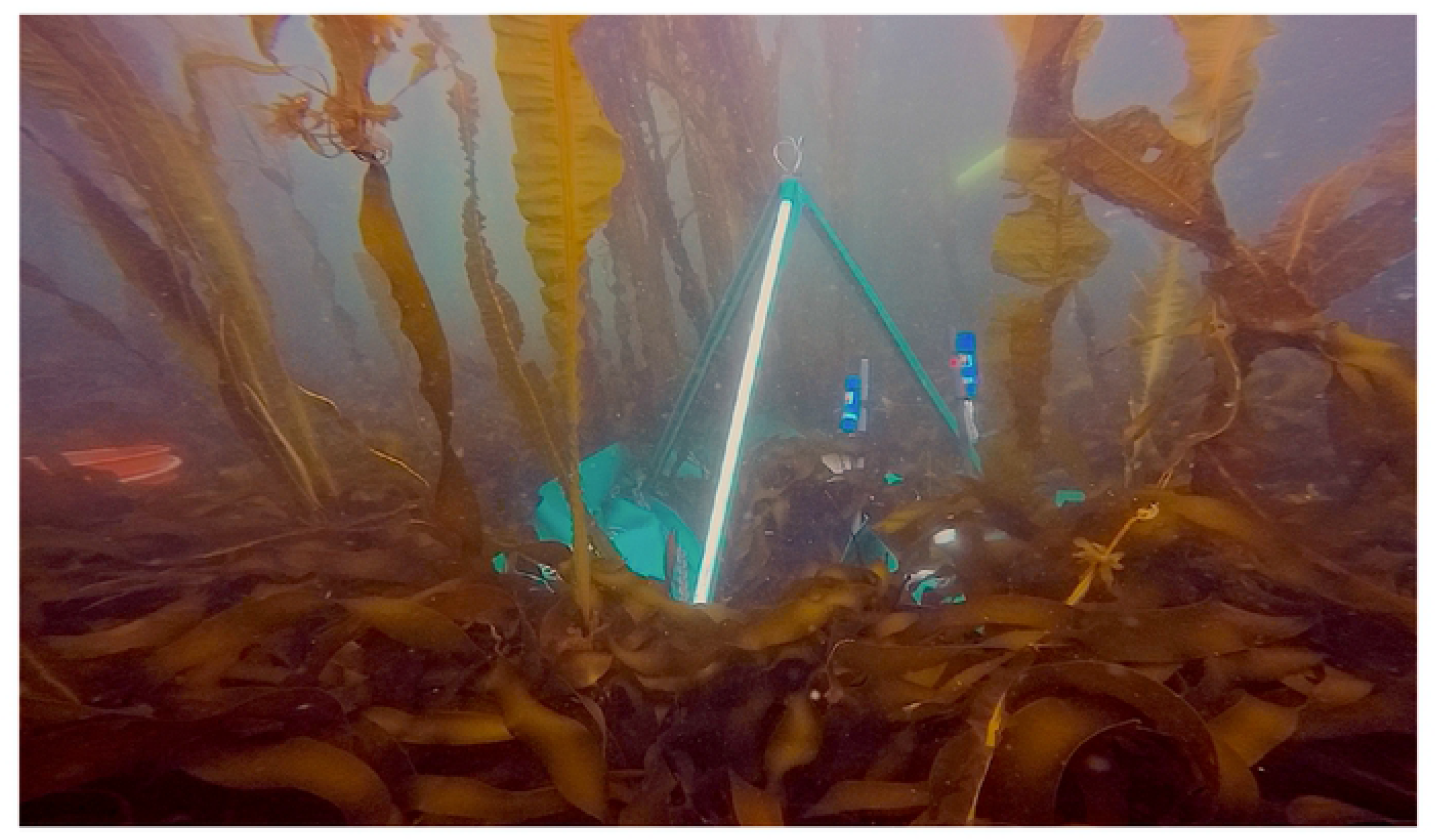
Photograph of cBIT deployed in kelp forest showing *PAR* and oxygen sensors

During two cruises aboard the *RV Oceanus* in 2016 and 2017, we deployed three cBITs in each of the three habitats (kelp forest, urchin barrens, transition habitats) on each of nine islands (Figs 2, 8) for 24-36 hour periods to measure both day and night patterns of *NEP* and *Re*. However, occasionally, replicates were lost due to logistical difficulties associated with the chamber-benthos seals (Table 1). For each deployment, the cBITs were placed over haphazardly-selected targeted assemblages in the field. The water within each cBIT was replaced once per day by opening the side of the chamber and completely replacing the water with new ambient seawater to reduce “chamber effects” (i.e. the build-up of oxygen and depletion of inorganic carbon and nutrients). After each deployment, the chambers and sensors were retrieved. At six of the islands (Table 1), all organisms within each of the chambers’ benthic footprints were collected, brought back to the ship, enumerated and weighed. We measured *NEP* over the whole diurnal cycle, *Re* during the nighttime hours, and calculated *GPP* during the day for each cBIT during each incubation period separately according to Olivé et al. (*46*). Specifically, measurements made during the night (the dark) were used to infer rates of *Re*, which were then combined with measurements of *NEP* to estimate *GPP* by the autotrophs (*19–21*).

### Statistical Analyses

All analyses were done in either Systat ver. 12, Primer ver 6. Prior to analyses, all data were evaluated for normality by graphical examination of the residuals, which suggested they were slightly non-normal. Data were then square-root transformed and re-graphed, which suggested the problems were corrected, with the exception of macroalgal biomass, which could not be fixed by transformation. The transformed data were then examined for equality of variances using Bartlette’s tests, which indicated they were homoscedastic. We then evaluated if urchin biomass, PAR, *GPP*, *Re*, *NEP* and the range between *GPP* and *Re* varied among the three habitats (kelp forests, urchin barrens, and transition habitats), and among islands using separate two-way Model III ANOVAs, with habitat type as a fixed factor, and island as a random factor.

Regardless of ANOVA outcomes, we then used Fisher’s LSD tests to evaluate *a priori* hypotheses about how these metrics differed between pairs of habitat types. We examined if macroalgal biomass varied among the three habitat types using Euclidean distance based PERMANOVA. Regardless of PERMNOVA outcome, we used permutation post hoc tests to evaluate *a priori* hypotheses about how biomass differed among the three habitat types. We evaluated if the relationship between *GPP* and *Re* varied among habitats using ANCOVA, with *Re* as the response variable, *GPP* as the covariate, and habitat type as the categorical independent variable. We evaluated if the ratios in any of the habitats differed from 1.0 (i.e. *GPP* = *Re*) by assessing if 1.0 occurred within the 95% confidence intervals around their average values.

## Acknowledgements

We thank S. Lamerdin, and the captain and crew of the *R/V Oceanus* for excellent ship support. We thank J. Estes for offering historical perspectives on the Aleutian kelp ecosystem, and M. Hatay for designing the cBITs. We are grateful to M. Good, S. Traiger, J. Metzger, A. Bland, A. Ravelo, and B. Weitzman for assistance with field operations. We also thank the Alaska Maritime National Wildlife Refuge for logistical support.

## Data and materials availability

Data are available on our NSF bco-dmo data page at https://www.bco-dmo.org/dataset/755658

## References

1. J. Terborgh, J. A. Estes, Trophic Cascades: Predators, prey and the Changing Dynamics of Nature. Island Press; 2010.

2. Ellison AM, Bank MS, Barton DC, E. A. Coulburn, Elliott K, et al. Loss of foundation species: consequences for the structure and dynamics of forested ecosystems. Front Ecol Environ. 2005; 3:479–486.

3. Ripple WJ, Becshta RL. Hardwood tree decline following large carnivore loss on the Great Plains, USA. Front Ecol Environ. 2004; 5:241–246.

4. Ripple, WJ. Wolves and the ecology of fear: can predation risk structure ecosystems. BioScience 2004; 54:55–766.

5. Beschta RL, Ripple WJ. Recovering riparian plant communities with wolves in northern Yellowstone, USA. Rest Ecol. 2010; 18:380-389.

6. Miller RJ, Harrer S, Reed DC. Addition of species abundance and performance predicts community primary production in macroalgae. Oecologia 2012; 168:797–806.

7. Wilmers CC, Estes JA, Edwards MS, Laidre KL, Konar B. Do trophic cascades affect the storage and flux of atmospheric carbon? An analysis of sea otters and kelp forests. Front Ecol Environ. 2012; 10:409–415.

8. Hondolero D, Edwards MS. Physical and biological characteristics of kelp forests in Kachemak Bay, Alaska. Mar Biol. 2017; 164:81–93.

9. Teagle H, Hawkins SJ, Moore P, Smale DA, The role of kelp species as benthic habitat formers in coastal marine ecosystems. J Exp Mar Biol Ecol. 2017; 492:81–98.

10. Krumhansl KA, Okamoto DK, Rassweiler A, Novak M, Bolton JJ, et al. Global patterns of kelp forest change over the past half-century. Proc Nat Acad Sci. 2016; 113:13785 – 13790.

11. Wernberg T, Krumhansl K, Filbee-Dexter K, Pedersen MF. “Chapter 3 – Status and trends of the world’s kelp forests” In World Seas: An Environmental Evaluation. Volume III: Ecological Issue and Environmental Impacts. 2^nd^ed. Academic Press. 2019, pp 57–78.

12. Estes JA, Tinker MT, Williams TM, Doak DF. Killer whale predation on sea otters linking coastal with oceanic ecosystems. 1998; Science 282:473-476.

13. Doroff AM, Estes JA, Tinker TM, Burn DM, Evans TJ. Sea otter population declines in the Aleutian archipelago. J Mamm. 2003; 84:55-64.

14. Edwards MS, Konar BK. A comparison of Dragon kelp, *Eualaria fistulosa*, (Phaeophyceae) fecundity in urchin barrens and nearby kelp beds throughout the Aleutian Archipelago. J Phycol. 2012; 48:897–901.

15. Konar BK, Edwards MS, Estes JA. Biological interactions maintain the boundaries between kelp forests and urchin barrens in the Aleutian Archipelago. 2014; Hydrobiol 724:91-107.

16. Costanza R, Fisher B, Mulder K, Liu S. Biodiversity and ecosystem services: A multi-scale empirical study of the relationship between species richness and net primary production. Ecol Econ. 2007; 61:478–491.

17. Harrisona PA, Berrya PM, Simpsona G, Haslettb JR, Blicharskac M, et al. Linkages between biodiversity attributes and ecosystem services: A systematic review Ecosyst Serv. 2014; 9:191–203.

18. Williams PJ, Purdie D. A. *In vitro* and *in situ* derived rates of gross production, net community production and respiration of oxygen in the oligotrophic subtropical gyre of the North Pacific Ocean. Deep-Sea Res. I. 1991; 38:891–910.

19. del Giorgio PA, Cole JJ. Photosynthesis or planktonic respiration? Nature. 1997; 388:132–133.

20. del Giorgio PA, Cole JJ, Cimbleris A. Respiration rates in bacteria exceed phytoplankton, Nature, 1997; 385:148–151.

21. Williams PJ. The balance of plankton respiration and photosynthesis in the open oceans. Nature. 1998; 394:55–57.

22. Duarte CM, Agusti S. The CO2 balance of unproductive aquatic ecosystems. Science. 1998; 281:234–236.

23. Staufenberger T, Thiel V, Wiese J, Imhoff JF. Phylogenetic analysis of bacteria associated with *Laminaria saccharina*. FEMS Microbiol Ecol. 2008; 64:65–77.

24. Minich JJ, Morris M, Brown M, Doane M, Edwards MS, et al. Elevated temperature drives kelp microbiome dysbiosis, while elevated carbon dioxide induces water microbiome disruption. 2018; PLOS ONE: PONE-D-17-36707R2.

25. Pfister CA, Alabet MA, Weigel BL. Kelp beds and their local effects on seawater chemistry, productivity, and microbial communities. Ecology. Forthcoming; https://doi.org/10.1002/ecy.2798

26. Middelboe AL, Sand-Jensen K, Binzer T. Highly predictable photosynthetic production in natural macroalgal communities from incoming and absorbed light. Oecologia. 2006; 150: 464–476.

27. Miller R, Reed DC, Brzezinski M. Community structure and productivity of subtidal turf and foliose algal assemblages Mar Ecol Prog Ser. 2009; 388:1-11.

28. Miller RJ, Reed DC, Brezinski MA. Partitioning of primary production among giant kelp (*Macrocystis pyrifera*), understory macroalgae, and phytoplankton on a temperate reef. Limnol Oceanog. 2011; 56:119-132..

29. Reisewitz S, Estes JA, Simestad CA. Indirect food web interactions: sea otters and kelp forest fishes in the Aleutian archipelago. Oecologia. 2006; 146:623–31.

30. Metzger JR, Konar B, Edwards MS. Assessing a macroalgal foundation species: community variation with shifting algal assemblages. Mar Biol. Forthcoming.

31. Teagle H, Hawkins SJ, Moore P, Smale DA. The role of kelp species as benthic habitat formers in coastal marine ecosystems. J Exp Mar Biol Ecol. 2017; 492:81–98.

32. Shukla J, Nobre Sellers CP. Amazon deforestation and climate change. Science. 1990; 247:1322–1325.

33. Brook BW, Sodhi, NS, Ng PK. Catastrophic extinctions follow deforestation in Singapore. Nature. 2003; 424:420–423.

34. Pfister CA, Berry HD, Mumford T. The dynamics of Kelp Forests in the Northeast Pacific Ocean and the relationship with environmental drivers. J Ecol. 2018; 106:1520–1533.

35. Filbee-Dexter K, Feehan CJ, Scheibling EE. Large-scale degradation of a kelp ecosystem in an ocean warming hotspot. Mar Ecol Prog Ser. 2016; 543:141– 152.

36. Raybaud V, Beaugrand G, Goberville E, Delebecq G, Destombe C, et al. Decline in kelp in West Europe and Climate. PLoS One https://doi.org/10.1371/journal.pone.0066044 (2013).

37. Tanaka K, Taino S, Haraguchi H, Prendergast G, Hiroka M. Warming off southwestern Japan linked to distributional shifts of subtidal canopy-forming seaweeds. Ecol Evol. 2012; 2:854–2865.

38. Jeon BH, Yang KM, Kim JH. Changes in macroalgal assemblages with sea urchin density on the east coast of South Korea. Algae. 2015; 30:139-146.

39. Martínez B, Radford B, Mads S, Thomsen, SD. Connell SD, et al. Distribution models predict large contractions of habitat-forming seaweeds in response to ocean warming. Divers Dist. 2018; 24:1350-1366.

40. Schiebling RE, Hennigar AW, Balch T. Destructive grazing, epiphytism, and disease: the dynamics of sea urchin – kelp interactions in Nova Scotia. Can J Fish Sci Aqu. 1999; 56:2300–2314.

41. Fagerli CW, Norderhaug KM, Christie HC. Lack of sea urchin settlement may explain kelp forest recovery in overgrazed areas in Norway. Mar Ecol Prog Ser. 2012; 488:119–132.

42. Bolton JJ, Anderson RJ, Smit AJ, Rothman MD. South African kelp moving eastwards: the discovery of *Ecklonia maxima* (Osbeck) Papenfuss at DeHoop Nature Reserve on the south coast of South Africa. African J Mar Sci. 2012; 34:147–151.

43. Edwards MS, Hernández-Carmona G. Delayed recovery of giant kelp near its southern range limit in the North Pacific following El Niño. Mar Biol. 2005; 147:273–279.

44. Tait L, Schiel D. Legacy effects of canopy disturbance on ecosystem functioning in macroalgal assemblages. PLOS ONE. 2011; 6:e26986.

45. Rodgers K, Rees T, Shears N. A novel system for measuring in situ rates of photosynthesis and respiration of kelp. Mar Ecol Prog Ser. 2015; 528:101–115.

46. Olivé I, Silva J, Costa M, Santos R. Estimating seagrass community metabolism using benthic chambers: the effect of incubation time. Estuar Coasts. 2016; 39:138–144.

47. Haas A, Nelson C, Rohwer F, Wegley-Kelly L, Quistad S, et al. Influence of coral and algal exudates on microbially mediated reef metabolism PeerJ. 2013; 1:e108.

48. Calhoun S, Haas A, Takeshita Y, Johnson M, Fox M, et al. Exploring the occurrence of and explanations for nighttime spikes in dissolved oxygen across coral reef environments. PeerJ Preprints. 2017; 5:e2935v1.

49. Glud R. Oxygen dynamics of marine sediments. Mar Biol Res. 2008; 4:243-289.

50. Bracken ME, Williams SL. Realistic changes in seaweed biodiversity affect multiple ecosystem functions on a rocky shore. Ecology. 2013; 94:1944–1954.

